# Optimizing murine sample sizes for RNA-seq studies revealed from large-scale comparative analysis

**DOI:** 10.1101/2024.07.08.602525

**Authors:** Gabor Halasz, Jennifer Schmahl, Nicole Negron, Min Ni, Gurinder S. Atwal, David J. Glass

## Abstract

Determining the appropriate sample size (N) for comparative biological experiments is critical for obtaining reliable results. In order to determine the N, the usual approach is to perform a power calculation, which involves understanding the variability between samples and the expected effect size. Here, we focused on bulk RNA-seq experiments, which have become ubiquitous in biology, but which have many unknown or difficult to estimate parameters, and so the required analyses to determine the minimum N is typically lacking. We therefore performed two N=30 profiling studies between wild-type mice and mice in which one copy of a gene had been deleted, to determine how many mice would be required to minimize false positives and to maximize true discoveries found in the N of 30 experiment. Results from experiments with N=4 or less are shown to be highly misleading, given the substantial false positive rate, and the lack of discovery of genes later found with higher N. For a cut-off of 2-fold expression differences, we found that an N of 6-7 mice was required to consistently decrease the false positive rate to below 50%, and that “more is always better” when it came to discovery rates - an N of 8-12 is significantly better in lowering the false positive rate.

A common method to reduce false discovery rate in underpowered experiments is to raise the fold cutoff or increase the stringency of the P-value and include only highly perturbed, highly significant genes. We show that while this strategy is no substitute for increasing the N of the experiment, because it results in consistently inflated effect sizes and a substantial drop in sensitivity. These data should be helpful to others in choosing their Ns, since it’s often not practical to do such large studies for every mouse model.

## INTRODUCTION

A typical RNA-expression study aims to find genes whose RNA levels differ significantly between two conditions. RNA-seq measurements are subject to technical noise and biological variability, the impact of which is expected to diminish with increasing sample size, N, in each group. Too few samples result in differentially expressed genes being missed (type 2 errors, or false negatives), spurious findings (type 1 errors, or false positives), and inflated effect sizes (type M errors, or the “winner’s curse” ^1, 2^). Underpowered studies are a major factor driving the lack of reproducibility in the scientific literature^3, 4^.

The intuitive preference for more replicates in study design is balanced in practice by resource constraints, ethical qualms about using more animals than needed, and by the career pressure on scientists to rapidly produce striking and publishable results. Unfortunately, this latter goal is widely achieved by statistically underpowered studies, compromising reproducibility, wasting the valuable resources and animals, and ultimately impeding scientific progress. Therefore, it is imperative to objectively assess effects of differing sample sizes and determine optimal guidelines for future experiments, reflecting an optimal tradeoff between the errors incurred at low N and the resource constraints at high N.

Analytical power calculations for RNA-seq studies are challenged by the observed long-tailed dispersed distribution of sequence count data, often modeled as a negative binomial distribution. Previous studies have employed parametric statistical tools that model power as a function of expected effect sizes, dispersion of the data, sequencing depth, and other factors^5–10^. Though potentially useful when accompanied by user-friendly tools, researchers seldom know the values of the parameters on which these models depend upon. More recent tools^10, 11^ estimate these parameters from existing studies that the researcher believes will be comparable to their planned one. Even so, different tools give discrepant results and perform poorly for low fold changes^12^.

These theoretical approaches are complemented by studies that infer appropriate sample sizes directly from empirical data. Baccarella, et al^13^ compare a human monocyte data set to a gold standard obtained from additional studies, finding that sample size has a much larger impact than read depth on precision and recall, with performance dropping notably below seven replicates. A similar analysis of six public data sets^14^ highlights the importance of sample size as well as gene expression variability (dispersion), and notes the challenge of accurately estimating the latter. Finally, S Church et al^15^ sub-sample from a large cohort of yeast to assess the impact of sample size on accurately calling differentially expressed genes, using their full cohort as the gold standard.

While these empirical studies highlight key principles of study design, none of them include the best studied model organism used in biology – the mouse. Mice are often studied due to the availability of inbred strains, which is hoped to decrease variability between study subjects; also, techniques for genetic manipulation – creation of heterozygous animals, knockouts, conditional knockouts, and transgenics – are well-established in mice. Finally, mice are mammals, and thus are more closely related to humans evolutionarily than other well-established genetic models such as yeast, C. elegans worms, drosophila, and zebrafish. In this study, we transcriptionally profile a large cohort of genetically modified and wild-type pure strain C57BL/6 mice, the most widely used mammal as preclinical models of human disease, and demonstrate that genotype comparisons between inadequately sized subsets of this cohort fail to recapitulate the full signature, and systematically overstate effect sizes. The appropriate adequately sized subsets was found to be a larger N than is often encountered in the literature, typically in the range of 3 to 6, casting doubt on the reported claims of differentially expressed genes, and thus setting new guidelines for future bulk RNA-seq experiments.

## RESULTS

In order to determine an ideal range of N for an RNA-seq study, we first performed a large scale set of comparative expression studies, with a maximum N=30, across four organs (heart, kidney, liver, and lung) from wild-type and heterozygous mice - in which one copy of a gene was deleted. This choice of maximum N, close to an order of magnitude larger than typically reported in published studies, was defined to be the gold-standard, capturing the true underlying biological effects as accurately as possible, and serving as a benchmark for comparison against subsets with smaller N. To mitigate the possible concern that a particular gene deletion may be untypically variable in gene expression changes across all tissues, we separately studied two distinct gene heterozygotes, resulting in a total of 360 RNA-seq samples. We sequenced 30 mice heterozygous for Dachsous Cadherin-Related 1 (Dchs1), 30 mice heterozygous for Fat Atypical Cadherin 4 (Fat4), together with 30 wild type (WT) mice, each group derived from the same litters; these heterozygous lines were picked as representative comparators versus wild-type animals. Dchs1 and Fat4 are large cell adhesion molecules that act as a tethered ligand–receptor pair on adjacent cells to mediate planar cell polarity. Homozygous null mutations of either gene in mice are lethal at neonatal stages and have similar phenotypes affecting many organs. These include postnatal lethality, decrease in body weight, small cystic kidneys, abnormal skeletal morphology, curly tails, small lungs and cardiovascular abnormalities. Heterozygous Dchs1 and Fat4 mice exhibit less severe phenotypes^16^. Every effort was made to control for confounding factors and reduce variation between individuals, including use of a highly inbred pure strain C57BL/6NTac line, identical diet and housing, IVF derivation from the same male, same day tissue harvesting and same day sequencing. We chose heterozygotes because we wanted to profile a system wherea particular gene was in place, but a phenotype was observable, as in the case of these animals - because many biological settings where RNAseq profile this sort of setting. In this text we focus our observations around Dchs1. Results for the kidney and liver of Fat4, included in the supplement, showed analogous patterns, while heart and lung yielded too few gene changes (12 and 7, respectively, see Supplementary Table 1) to examine meaningfully.

Dchs1 heterozygous (Het) mice showed strong gene expression changes relative to WT mice in all four tissues assayed. The liver and kidney showed the most perturbations, with key tissue markers and functions strongly affected. Gene signatures derived using the full 60 (30 versus 30 comparison) mouse cohort are designated the gold standard for differentially expressed genes (DEGs). Note that a separate gold standard set is calculated for each combination of P-value, fold change, and absolute expression thresholds considered.

We assessed the impact of replicate number on the sensitivity and false discovery rate (FDR) using a down-sampling strategy (Figure 1). For a given sample size N, we randomly sampled N Het and N WT samples without replacement (N ranges from 3 to 29), performed DEG analysis, and compared the resulting signature to the gold standard (N=30). We define sensitivity as the percent of gold standard genes detected in the sub-sampled signature, while genes found only in the latter are considered false discoveries. Figure 2A shows the results of these virtual experiments (40 Monte Carlo trials for each N), using a fold change cutoff of 1.5 (50% up- or down-regulation) and adjusted P-value of less than 0.05. As expected, FDR drops towards zero while sensitivity rises towards 100% as N increases and the experiments more closely resemble the gold standard one. The FDR appears to exhibit an elbow, or inflection point, at around N=10 to 12, depending on tissue, indicating diminishing returns at higher N values. Sensitivity increases more smoothly, after a marked jump from N=4 to 5. For heart, kidney, and liver, a sensitivity of 50% is attained by N=8, while the lung required a sample size of 12. The liver signature of Fat4 showed analogous results, with an FDR inflection point around N=8 (Figure S5). Sensitivity increased smoothly, though without the jump. Fat4 kidney did not have an obvious inflection point, and sensitivity jumped at a higher N∼9. Overall, false discovery rates were higher, and sensitivity lower, in the Fat 4 Hets than in Dchs1, reflecting the general tendency that a stronger overall effect (as reflected by the number of genes perturbed) leads to better agreement.

**Figure 1.**
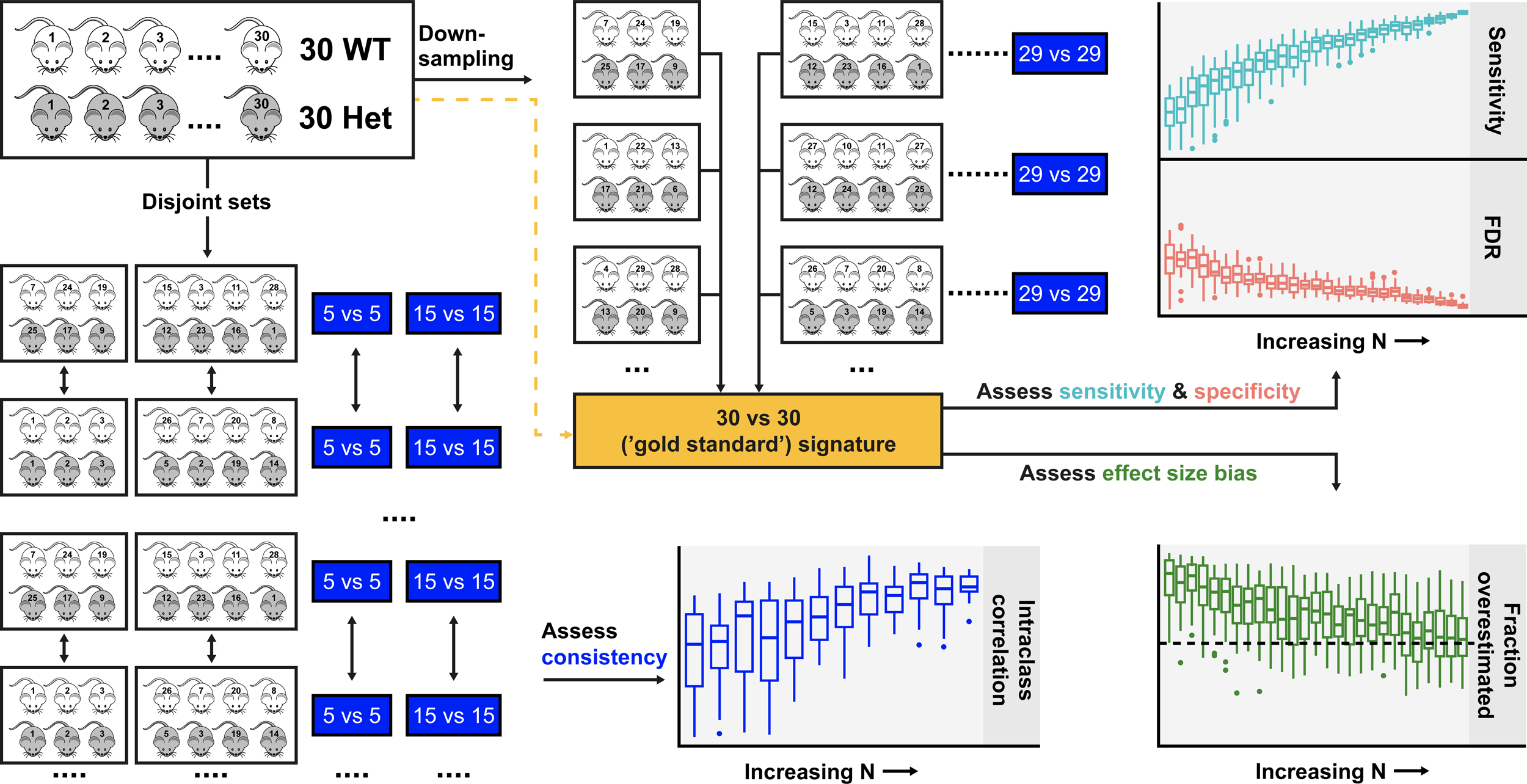
Study Overview. **Workflow for assessing the impact of sample size on discovery metrics.** We define the ‘gold standard’ signature as the set of DEGs in the full cohort of mice (30 Dchs1 Het vs 30 WT). From this full cohort, we generated a smaller “mini-experiment” (trial) by randomly selecting N Het and N WT mice (**Down-sampling** strategy, top middle). The DEG from this trial were compared to the gold standard signature to assess sensitivity, specificity, and effect size bias. The number of mice selected (N) range from 3 to 29, and 40 trials were performed for each N. A separate approach (**disjoint sets**) was used to assess consistency between pairs of mini-experiments. Here two mini-experiments of size N are created in each trial, and the resulting two DEG sets are compared. Higher consistency between DEGs is interpreted as each DEG capturing more biological signal.

**Figure 2.**
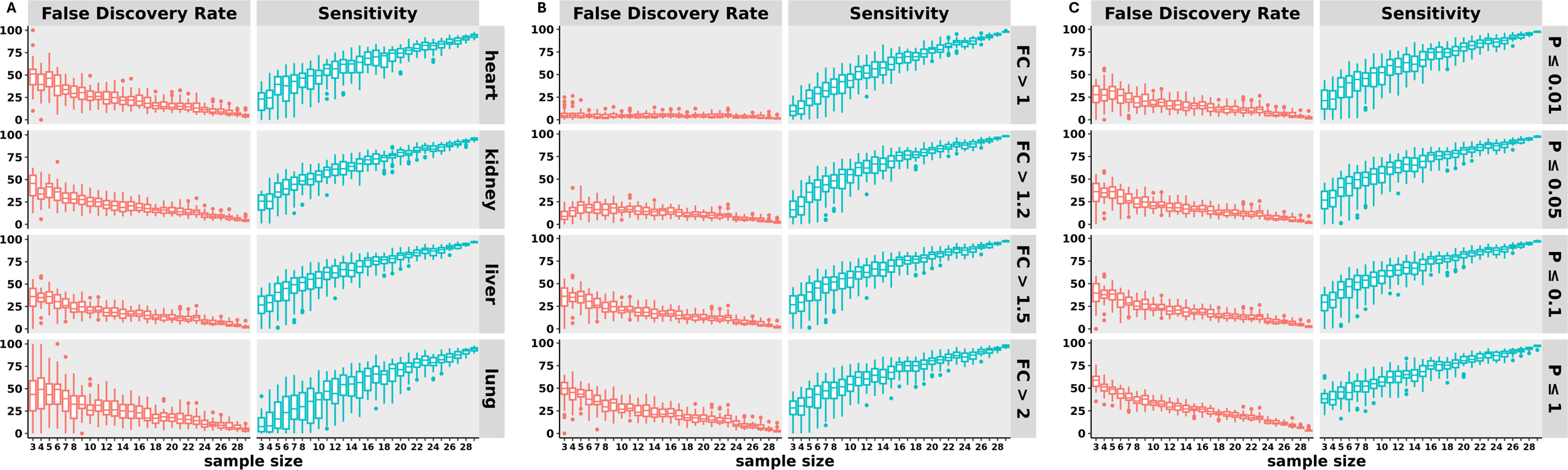
Sample size has major impact on sensitivity & false discovery rate for all tissues assayed. **A)** For each sample size N (x-axis), N Dchs1 Hets and N WTs were randomly chosen, Het-vs-WT DEGs calculated, and compared to the gold standard (full cohort) signature. Left panel shows FDR, calculated as the fraction of DEGs from the sub-sampled trial that was absent from the gold standard signature. Right panel shows sensitivity, defined as the fraction of gold standard DEGs also detected in the sub-sampled trial. Signature genes were those perturbed by 50% or more, with an adjusted P-value < 0.05. **B)** Plots are as in (A), except showing a single tissue, liver, with varying fold change thresholds along each row. C) Plots are as in (A), except showing a single tissue, liver, with varying P-value thresholds along each row.

The variability in false discovery rates across trials markedly drops with larger sample sizes. In the heart and lung, the FDR ranges between 10 and 100% depending on which N=3 mice are selected for each genotype. Some variability persists even with double-digit sample sizes, particularly in the Fat4 experiment. Notably, this variability is lower in kidney and liver, the tissues most affected by the genetic modification. The variation in sensitivity across trials also inversely varies with N for these highly impacted tissues, but this relationship is less apparent in lung, or in the Fat4 Hets.

We next examined the impact of varying fold change thresholds. Can researchers salvage an underpowered study by limiting their purview to highly perturbed genes? As shown in Figure 2B, narrowing the focus to high effect sizes increases the false discovery rate, as compared to a gold standard using the same fold change cutoff. Intuitively, this happens because achieving a statistically significant P-value in an underpowered experiment requires an extreme, and so likely inflated, effect size that was not observed in the gold standard signature. Note that for a given N, high-fold-change genes are more likely than low-fold-change ones to be perturbed at all, as shown in Supplementary Figure S2A. However, the observed fold change should not be taken at face value. In addition, sensitivity to detect true changes drops markedly, as we would expect (Supplementary Figure S2B). Increasing the stringency of the P-value, rather than fold change filter, yields similar, though slightly less pronounced, increases in FDR (Fig 2C). By contrast, applying a minimum absolute expression threshold does not lead to a similar FDR increase (Fig S1C). The above observations are corroborated in all tissues assayed (Fig S1A & S1B), and in the Fat4 experiment (FigS6A & S6B).

A limitation of the down-sampling approach for assessing an “optimal” sample size is that the same mice are used to derive the gold standard and the random trial gene signatures. Our second approach avoids this circularity by randomly sampling mice to create two disjoint experiments of size N in each trial, and comparing their DEGs (Figure 1, bottom). As N increases, signatures from the two sub-sampled, independent experiments should both better capture the underlying common biology and hence resemble each other more. Figure 3 shows that this holds for all tissues and fold change thresholds tested. Except for lung, the fraction overlap between DEGs reaches an asymptote at around N=12, in line with the inflection point observed with the down-sampling approach (Figure 2A). Fat4 liver showed a similar pattern, though with lower overall concordance. Interestingly, signatures from Fat4 kidney samples showed little concordance even with large N (Figure S7), suggesting that the initial gold standard signature may itself not be robust. As with the other approach, limiting to genes with a minimum absolute expression does not appreciably change these results (Figure S3).

**Figure 3.**
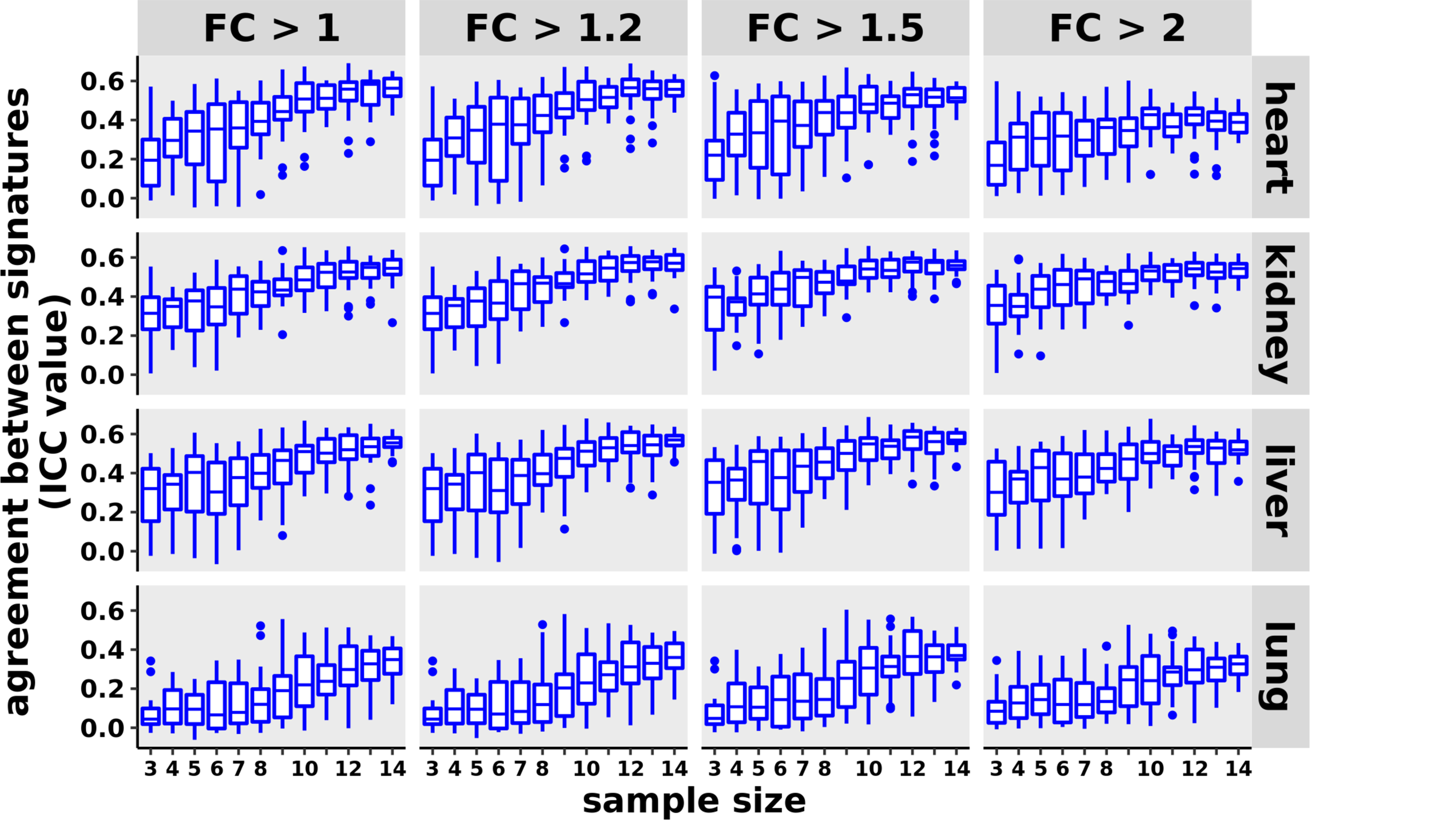
Agreement between DEGs obtained using disjoint method. The sample size N is shown along the x-axis, while the y-axis shows ICC calculated between expression signatures from two independently constructed experiments of size N. ICC values range from zero (no agreement) to one (perfect agreement, identical DEGs).

In addition to the loss of sensitivity and specificity, underpowered studies lead to inflated effect size estimates. To study this further, we compared the fold changes of DEGs shared between the gold standard and down-sampled trials. For small sample sizes, almost all DEGs overestimate the fold change in all tissues (Figure 4A), regardless of the cutoff for fold change or absolute expression. Even at N=12, considerably greater than the expected 50% of DEGs show perturbations more extreme than those observed in the gold standard signature. Fat4 recapitulates this pattern (Fig S8).

**Figure 4.**
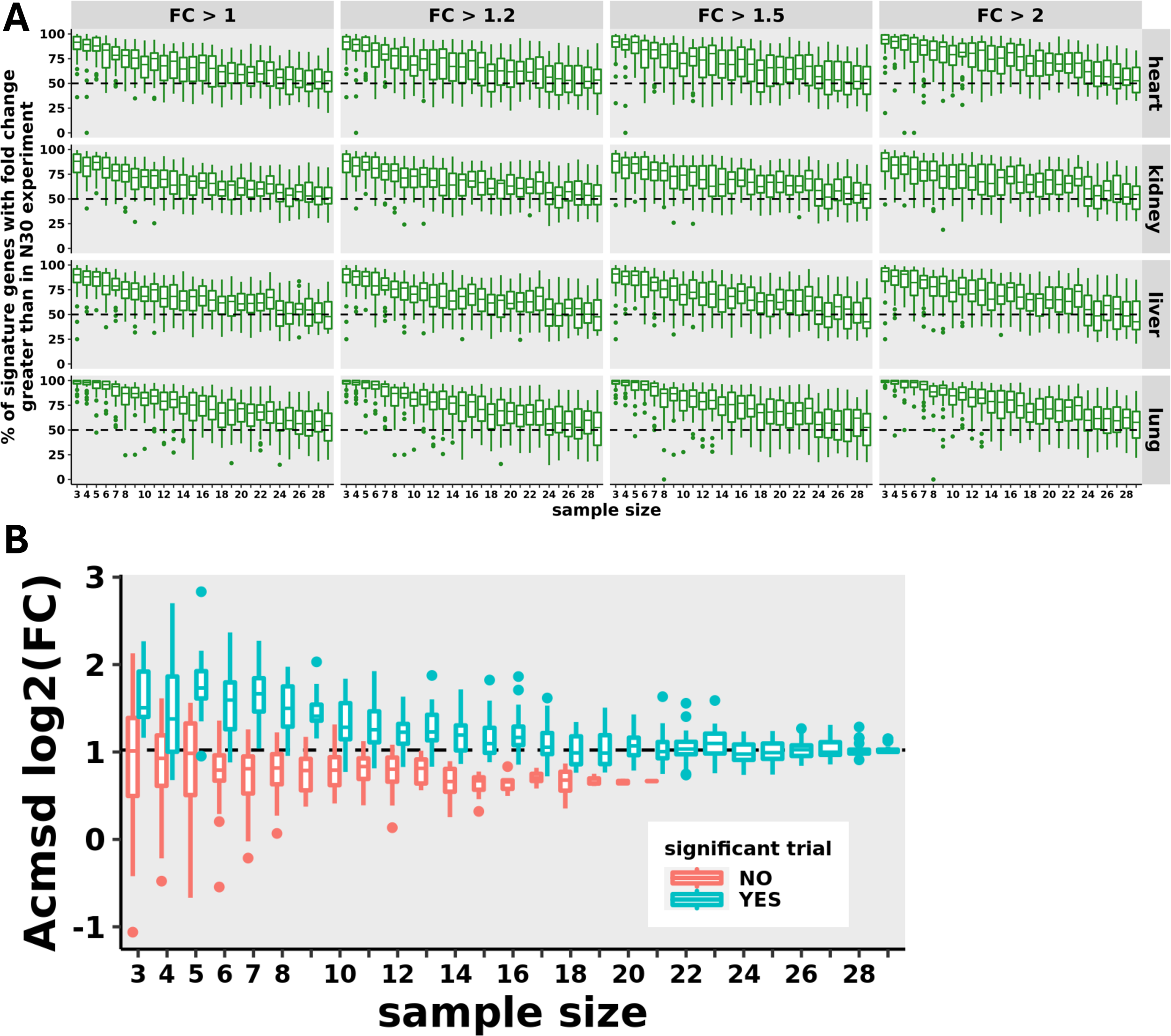
Impact of sample size on estimated effect size. **A)** Plots show the percentage of true positive genes in each trial whose absolute fold change in the sub-sampled DEG exceeds that of the fold change in the gold standard DEG. Sample size N is shown along the x-axis. The dotted line at 50% indicates no bias, where a gene is equally likely to over- or under-estimate the effect. B) Effect size estimates for a representative gene (Acmd) in liver. For each N (x-axis), the Het-vs-WT fold change estimate of Acmd is shown separately for statistically significant (cyan), or not significant (red) trials. Statistical significance was defined as having a BH-adjusted P-value of 0.05 or lower. With small N, fold change estimates from significant trials overestimate that of the gold standard (solid horizontal line), but approach it as N increases. Non-significant trials by contrast show no bias at small N.

Focusing on a representative example, Mkln1 in liver, we observe this fold change inflation relative to the gold standard (horizontal black line in Figure 4B) – but only among trials yielding a statistically significant P-value. In trials where the gene expression change did not reach significance, the fold change is accurately estimated, though with considerable variation around this estimate. The P-value filter thus acts as an effect overestimate generator, since only large effect sizes attain significance in underpowered trials. With increasing N and adequate power, the observations reverse: estimated fold changes for significant trials approach the gold standard estimate, while non-significant trials deviate from it. With high enough N, non-significant trials disappear altogether. Figure S4 shows several randomly chosen gold standard genes for each tissue, demonstrating the generality of these observations.

## DISCUSSION

Over the last several years, there has been an increased focus on “rigor and reproducibility” when it comes to biological data. This is a result of several studies indicating that quite a few high profile papers were found to be non-reproducible upon attempts to repeat those studies^17, 18^. What followed were many discussions as to what might be improved – and suggestions ranged from focus on journal policies^19^, education^20–22^, and policies of funding organizations^23^.

Of course, a fundamental component to addressing reproducibility is to perform studies that directly query the ability to reproduce the data from a particular type of experiment. Bulk tissue RNA-seq studies have become an indispensable and widespread technology for transcriptome wide analysis of DEGs over the past 15 years, and remain a staple of current comparative functional genomics. Given this ubiquity, it is important to understand what range of N is required to better assure that the conclusions made will be found to be predictive of subsequent analyses of the same sort of comparison.

One particularly common bulk RNA-seq experiment involves comparing knockout or transgenic animals with wild-type controls. When a gene is deleted, it is of interest to learn how this change perturbs mRNA expression, because this gives indications as to the signaling pathways which are normally affected by the gene of interest. We chose heterozygotes in this study because we wanted to profile a system wherea particular gene was in place, but a phenotype was observable, as in the case of the animals used here - because many biological settings where RNAseq is performed profile this sort of setting. As a comparator, wild-type animals are used – animals in which the gene is left unperturbed.

A particularly common preclinical model for such experiments are c57bl6 mice. Since this is an inbred mouse strain, one might expect that genetic variation would be minimized in comparison to outbred animals; indeed, the ability to use fewer animals is a common rationale to study inbred strains of mice. Even though this inbred mouse line is relatively genetically homogenous in comparison to outbred, or truly “wild” mice, this study makes it clear that considerable variety in gene expression across mice still persists, with implications for study design and interpretation.

Our results demonstrate how underpowered RNA-Seq experiments result in type I, type II, and type M (magnitude) errors, and offer guidance about adequate sample sizes to mitigate them. Commonly reported sample sizes of 5 or less are to be avoided, since for our studies seeking to identify genes perturbed by at least 1.5-fold, one finds about 50% false discoveries, less than 50% power to detect true changes, and inflated fold changes nearly 100% of the time. Considerable discrepancies persist between the gold standard N=30 signature and those from sub-sampled experiments, which diminish with increasing N. So one simple conclusion is that “more is always better” in the case of sample size – there is no point at which adding samples doesn’t help reduce error. Given realistic constraints such as budget and colony sizes, we looked for a point of diminishing returns. Both the down-sampling and disjoint approach suggest that this is reached around an N of 8-12.

Nearly all observed fold changes overestimate the true effect for N less than 6-7, though considerable over-estimation persists even at higher N. The reason for this is that smaller, underpowered studies require a larger effect size to achieve significance. This observation also explains why false discovery rates increase as we raise the fold change cutoff for both the sub-sampled and gold standard experiments: the latter tend to show more modest changes, and so genes whose fold change falls below threshold will be counted as a false positive. The intuitive solution is to apply a stringent cutoff to only the low-N experiment, while keeping in mind that the true changes are almost certainly more muted than those observed. In Figure S2, we define the gold standard signature using a modest 1.5 fold change cutoff. To achieve a 50% false discovery rate against this signature among genes with observed fold change between 1.5 and 2 requires an N of 6 or 7, depending on the tissue. Sample size around 14 is required for a 25% false discovery rate. However, focusing on genes with fold change greater than 2 greatly reduces false discoveries, while genes with fold change greater than 3 are almost guaranteed to be at least modestly perturbed in the 30-vs-30 experiment. This increase in specificity is unfortunately offset by a substantial drop in sensitivity, our ability to detect true gene changes. The widespread practice of focusing on highly perturbed genes therefore fails to fully capture the biology of the studied model; higher sample sizes are needed.

We observed similar observations with two separate sets of heterozygous animals – analyzing two distinct genes. Given that these relatively high Ns were found for an inbred strain, it should also be acknowledged that it’s highly likely that even higher Ns would be needed for outbred strains, as well as human studies. One might ask whether there is some idiosyncracy in the two heterozygotic lines studied that would make the data adduced in this study more variable than that found in other settings. Of course, the only way to answer this would be to do similar studies in still more genetically modified lines vs wild-type controls. The expression and function of Dchs1 and Fat4 are not known to be circadian, or feeding-dependent, for example, or dependent on other highly variable factors. Therefore, there is no particular reason to believe that the two heterozygous strains highlighted in this paper are unusually variable.

We hope this study will be generally useful for the determination of N in future RNA-seq studies, and in evaluating the utility of prior-published work. This study should also help contextualize studies done with comparatively low Ns.

## Acknowledgments

We thank Drs. L.S. Schleifer, G.D. Yancopoulos and A. Murphy, along with the rest of the Regeneron Community for their support. We would also like to thank: the Regeneron DNA Core AND the Molecular Profiling Lab, including Joseph Song, Shuo Li, and Nicole Negron for their work on the RNA-Seq assay. All authors were employees of Regeneron when this study was conducted; some hold stock in the company.

## MATERIALS AND METHODS

### Transgenic mice

Samples were obtained from 11-12 week old C57BL/6NTac male mice after overnight fasting. As Fat4 and Dcsh1 are lethal as homozygotes, comparisons were done between heterozygous and WT mice taken from the same litters. Both the Fat4 and Dchs1 deletions removed cExon1, starting at the ATG. LacZ was used as a reporter

### RNASeq analysis

RNA was prepped from tissues stored in RNA*later* using MagMAX Nucleic Isolation Kits on KingFisher Instruments (ThermoFisher). Strand-specific RNA-seq libraries were prepared from 500 ng RNA using KAPA Stranded RNA-Seq Kit for Illumina Platforms (Roche). Twelve-cycle PCR was performed to amplify libraries. Sequencing was performed on Illumina HiSeq®2500 (Illumina) by multiplexed sequencing with 33 cycles.

Raw sequence data (BCL files) were converted to Fastq format via Illumina Casava 1.8.2. Reads were decoded based on their barcodes, and read quality was evaluated using Fastqc (www.bioinformatics.babraham.ac.uk/projects/fastqc/). Reads were mapped to the mouse transcriptome (NCBI genome assembly GRCm38/mm10) using ArrayStudio’s OSA aligner, allowing for two mismatches. Exon mapped reads were summed at the gene level.

Differentially expressed genes were obtained using DeSeq2^24^ (1.34.0), with default parameters. Down-sampling (figures 1B-D, 3) was performed by selecting N WT and N Het mice without replacement from the initial 30+30 cohort. DEGs were then derived for all four tissues assayed using the selected mice and compared to the gold standard (N=30) signature. For the alternative, “disjoint” approach (figure 2), two non-overlapping sets of N WT and N Het mice were selected, and the DEGs from these two sets were compared and quantified using the intra-class correlation coefficient (ICC). For showing effect size differences, representative genes were chosen randomly from each tissue’s gold standard DEG list, with adjusted P-value threshold of 0.05. For each tissue, 20 genes were selected, evenly distributed among: DEGs with absolute fold change between 1-1.2; between 1.2-1.5; between 1.5-2; and greater than 2.

## Supplementary Figure Legends, Figures and Table

**Supp. Table 1:** Number of significantly perturbed genes in gold standard (N=30-vs-30) experiments.

| Tissue | Dchs1-Het vs WT | Fat4-Het vs WT |
| --- | --- | --- |
| heart | 846 | **12** |
| kidney | 928 | 57 |
| liver | 1415 | 110 |
| lung | 681 | 7 |

**Supplementary Figure 1A.**
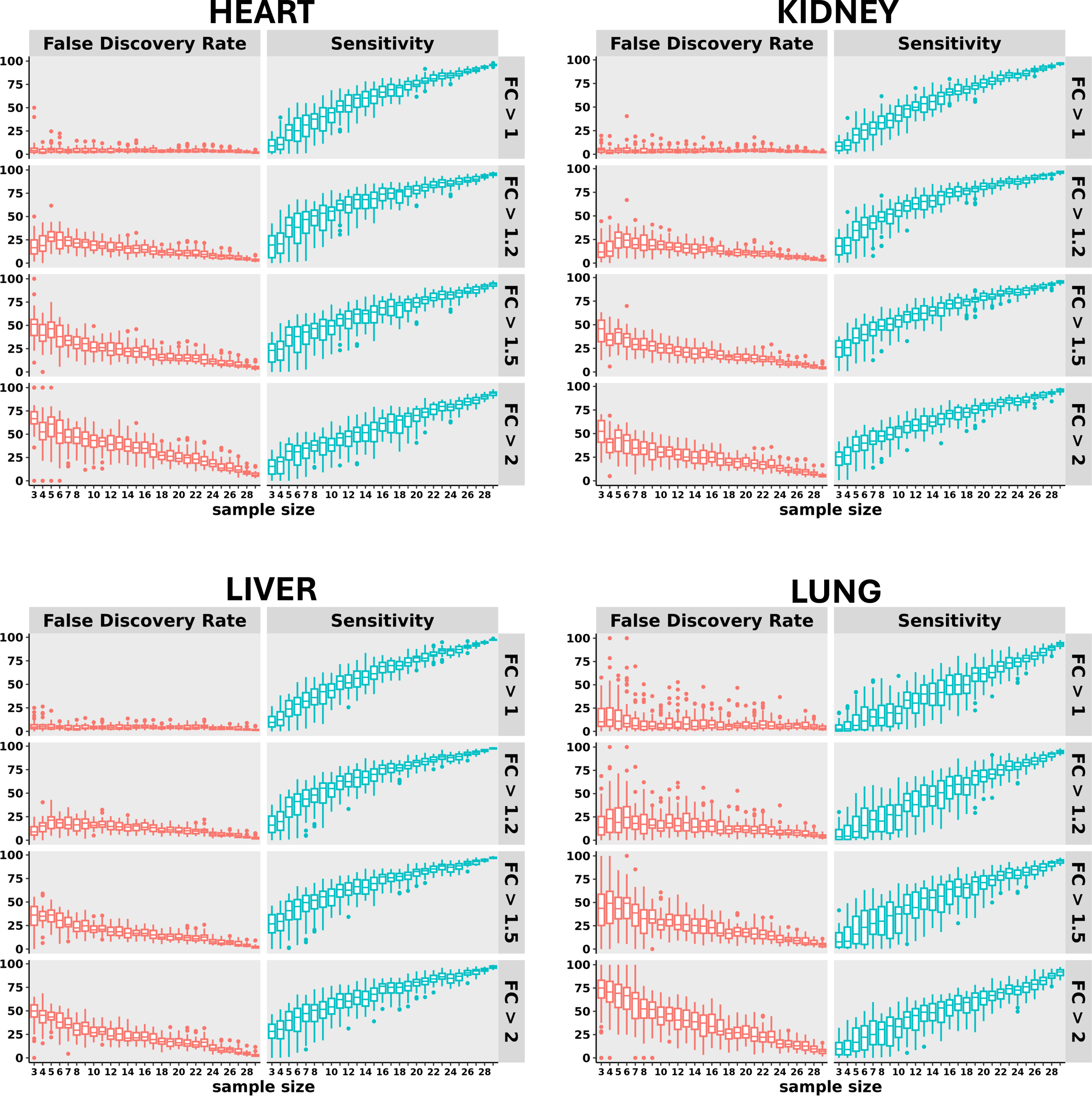
Impact of sample size and fold change threshold on sensitivity & false discovery rate in non-liver tissues of Dchs1 Het mice. Plots are constructed as in Figure 2B, but for all tissues assayed. Liver results (from Figure 2B) included here for reference.

**Supplementary Figure 1B.**
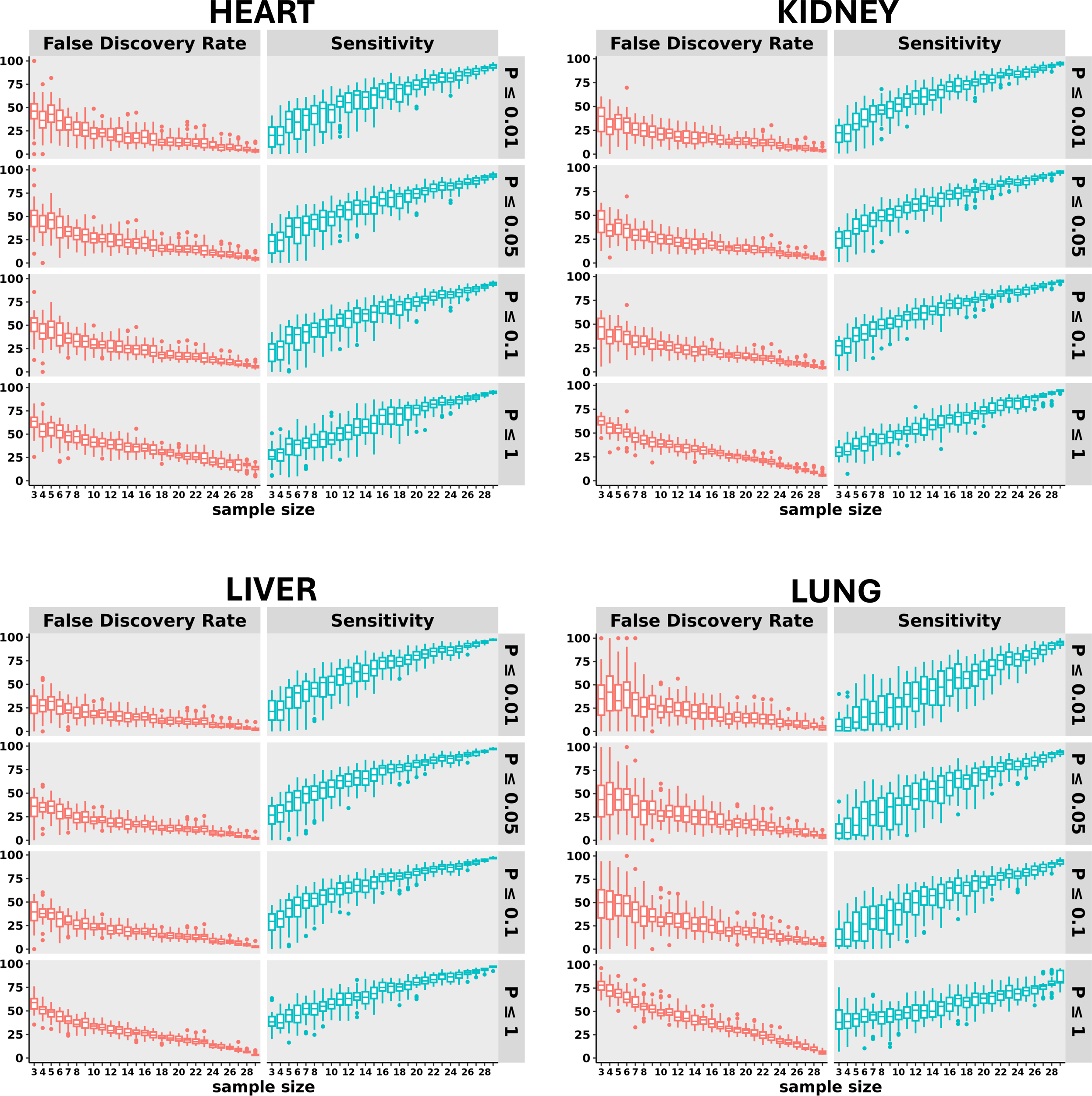
Impact of sample size and P-value threshold on sensitivity & false discovery rate in non-liver tissues of Dchs1 Het mice. Plots are constructed as in Figure 2C, but for all tissues assayed. For all panels, fold change cutoff is 1.5, and adjusted P-value thresholds are as shown. Liver results (from Figure 2C) included here for reference.

**Supplementary Figure 1C.**
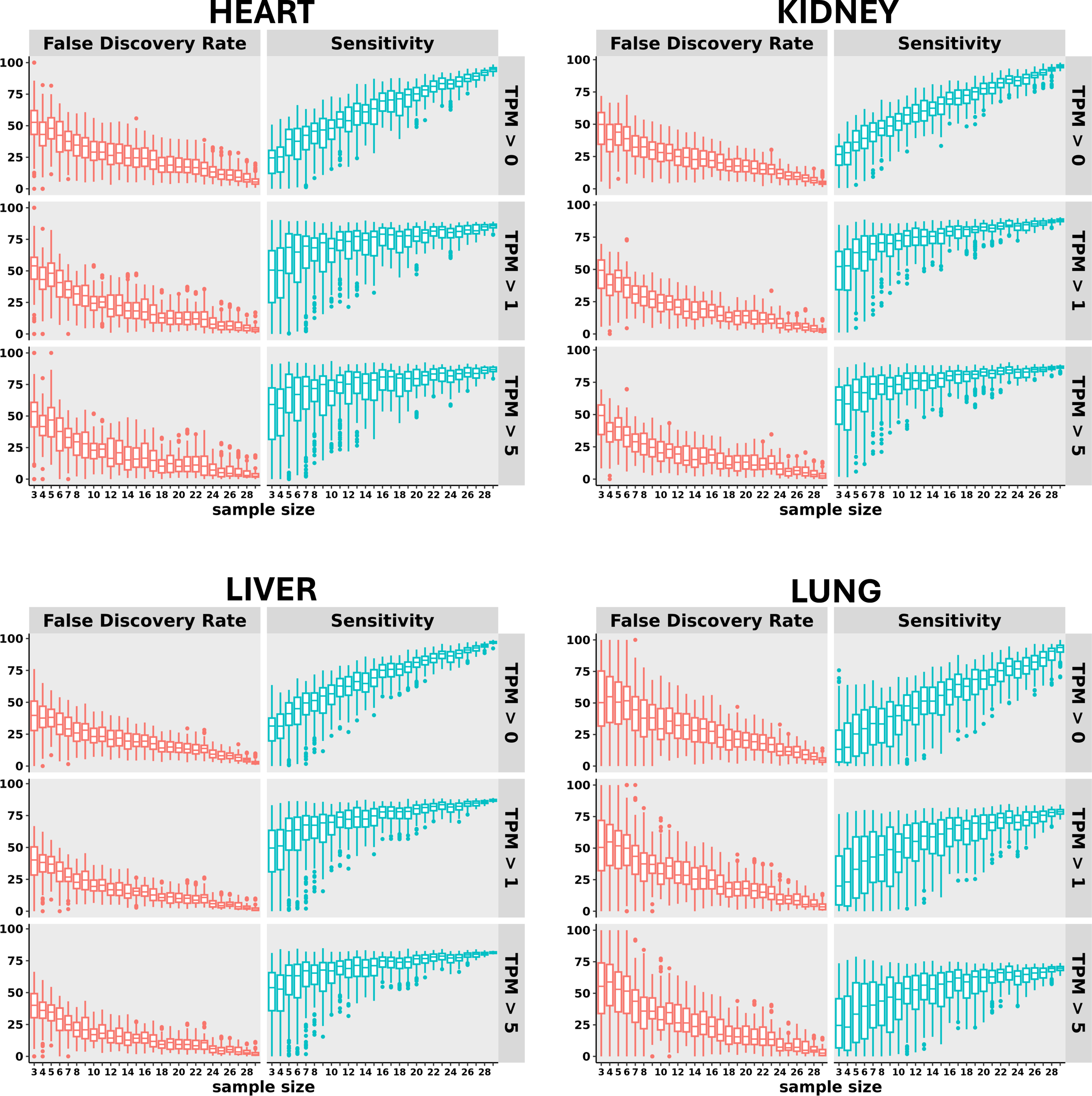
Impact of sample size and absolute expression threshold on sensitivity & false discovery rate in non-liver tissues of Dchs1 Het mice. Plots are constructed as in Figure 2B & 2C, but varying minimum expression (TPM). Genes were considered differentially expressed if they had an absolute fold change ≥ 1.5, adjusted P-value ≤ 0.05, and mean absolute expression in the higher-expressed arm of the comparison as indicated in the vertical panels.

**Supplementary Figure 2A.**
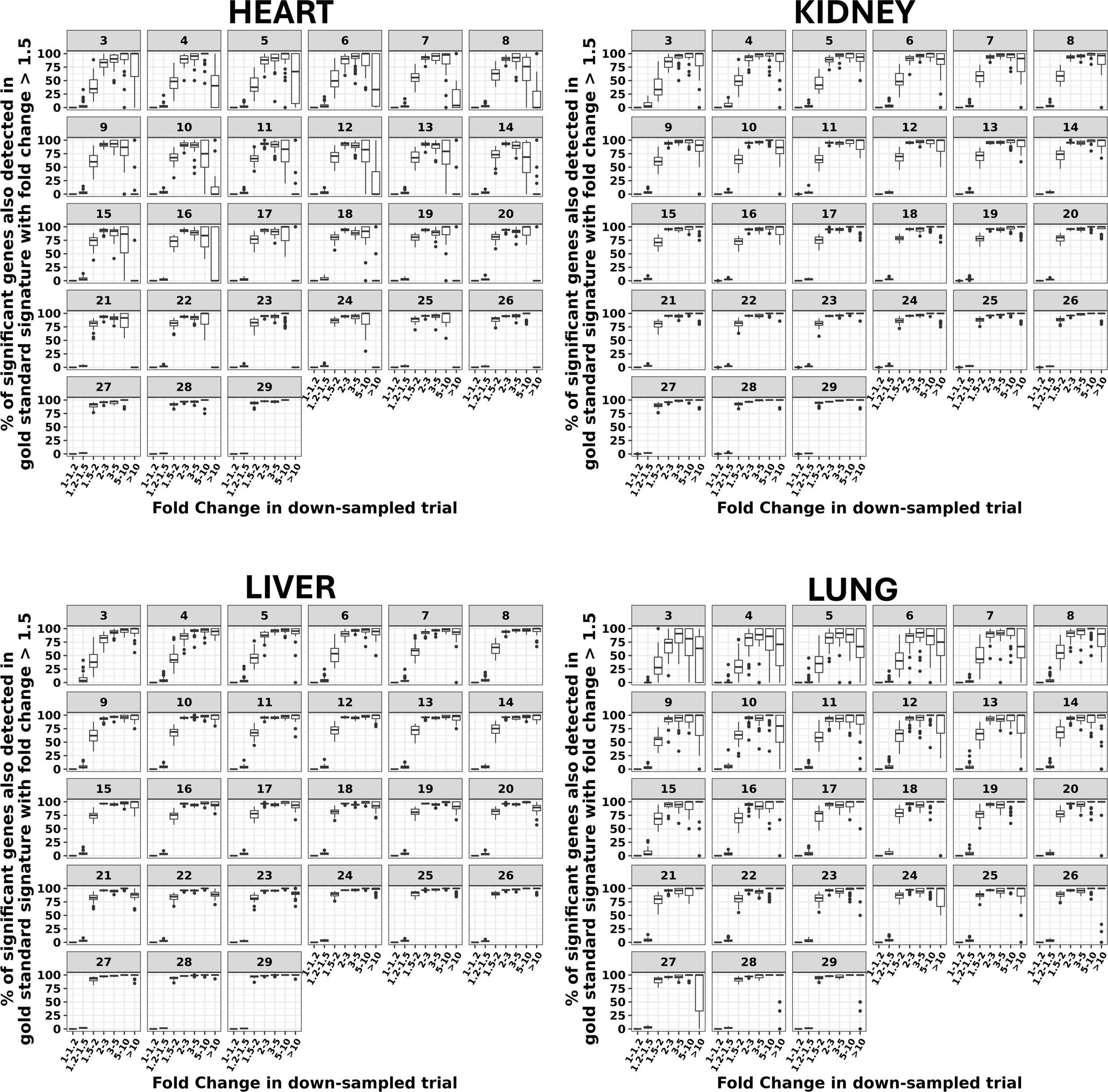
Genes with higher fold change in a down-sampled trial are more likely to be found in the gold standard signature. Likelihood of a gene perturbed in a down-sampled trial being found in the indicated tissue’s gold standard (GS) signature, for a fixed GS fold change threshold of 1.5. Significantly perturbed genes with fold change less than 1.5 in the trial are almost always false discoveries; those between 1.5-2 are usually false discoveries for N < 7, and don’t reach 80% likelihood of being in the GS signature unless very high N (>15) are used. Genes with fold change greater than 2 overwhelmingly represent true discoveries (for N > 5). However, as figures 2B and 4 suggest, these observed fold changes will be overestimates. (Interestingly, the trend towards increasing likelihood of true discovery reverses with very high (>10) fold change in the trial. These may be due to low overall expression leading by chance to highly inflated fold changes. Extreme fold changes should therefore be rigorously examined before being taken at face value).

**Supplementary Figure 2B.**
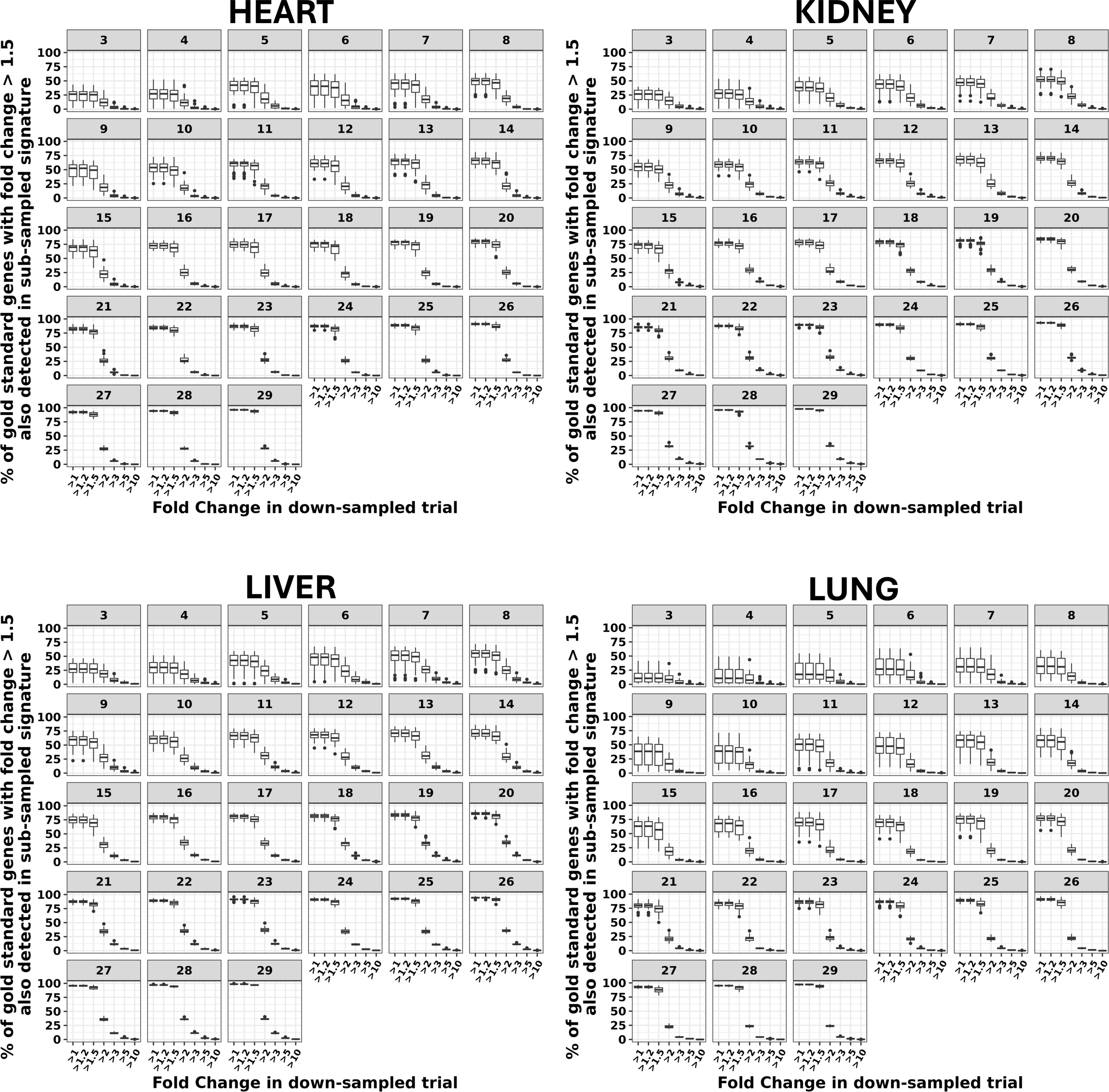
Increasing fold change stringency in down-sampled trial reduces sensitivity to detect changes in gold standard signature. Sensitivity of a down-sampled trial with varying fold change thresholds to detect genes in gold standard (GS) signature (GS fold change threshold is fixed at 1.5). For all tissues, sensitivity drops markedly if we increase the fold change threshold for the down-sampled trial from 1.5 to 2.

**Supplementary Figure 3.**
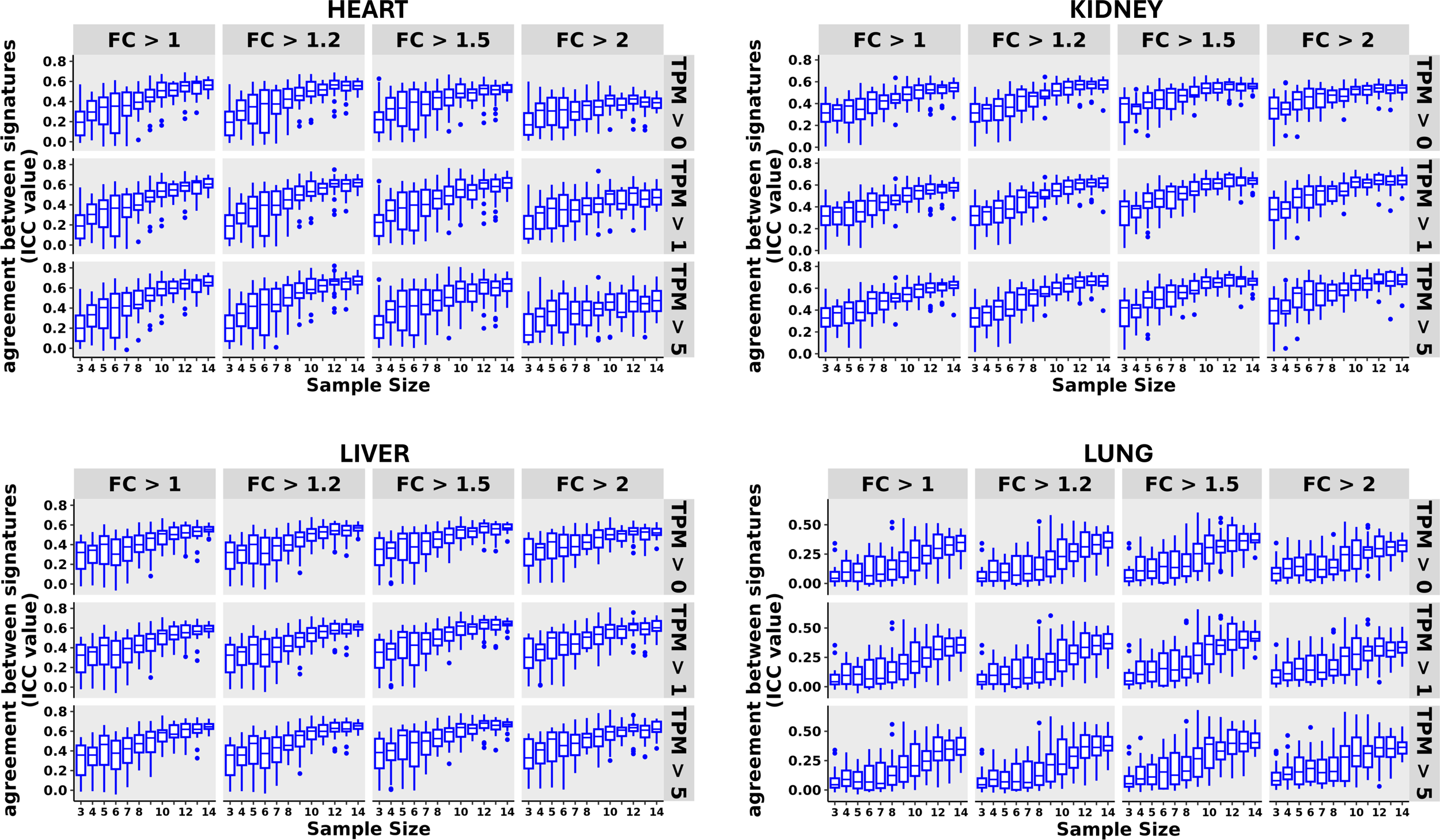
Impact of sample size and absolute expression threshold on DEG agreement using disjoint method. Plots constructed as in Figure 3, but showing the (modest) impact of imposing an absolute expression threshold on genes considered significantly perturbed.

**Supplementary Figure 4.**
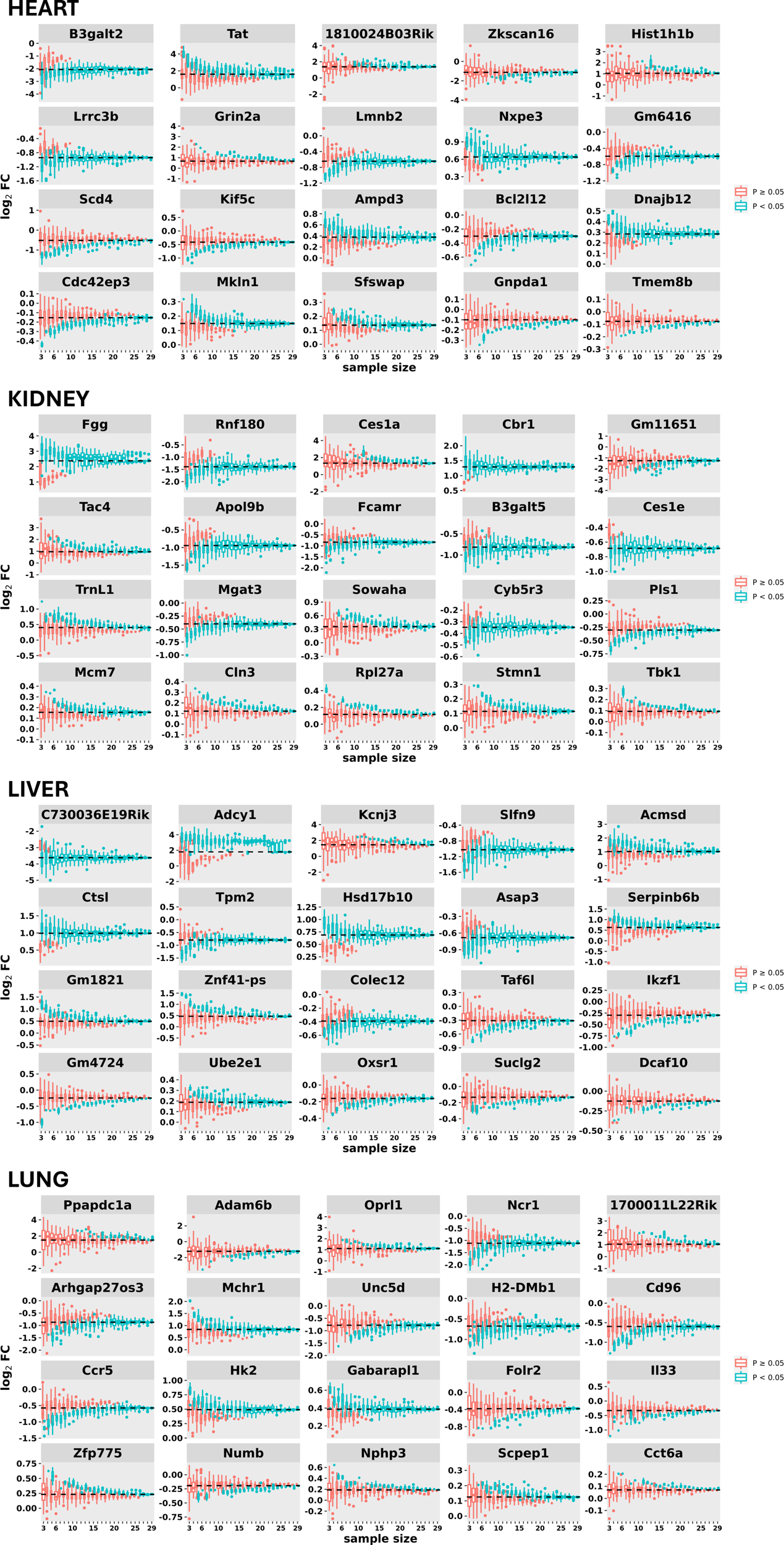
Additional randomly selected DEGs showing generality of effect size bias.

**Supplementary Figure 5.**
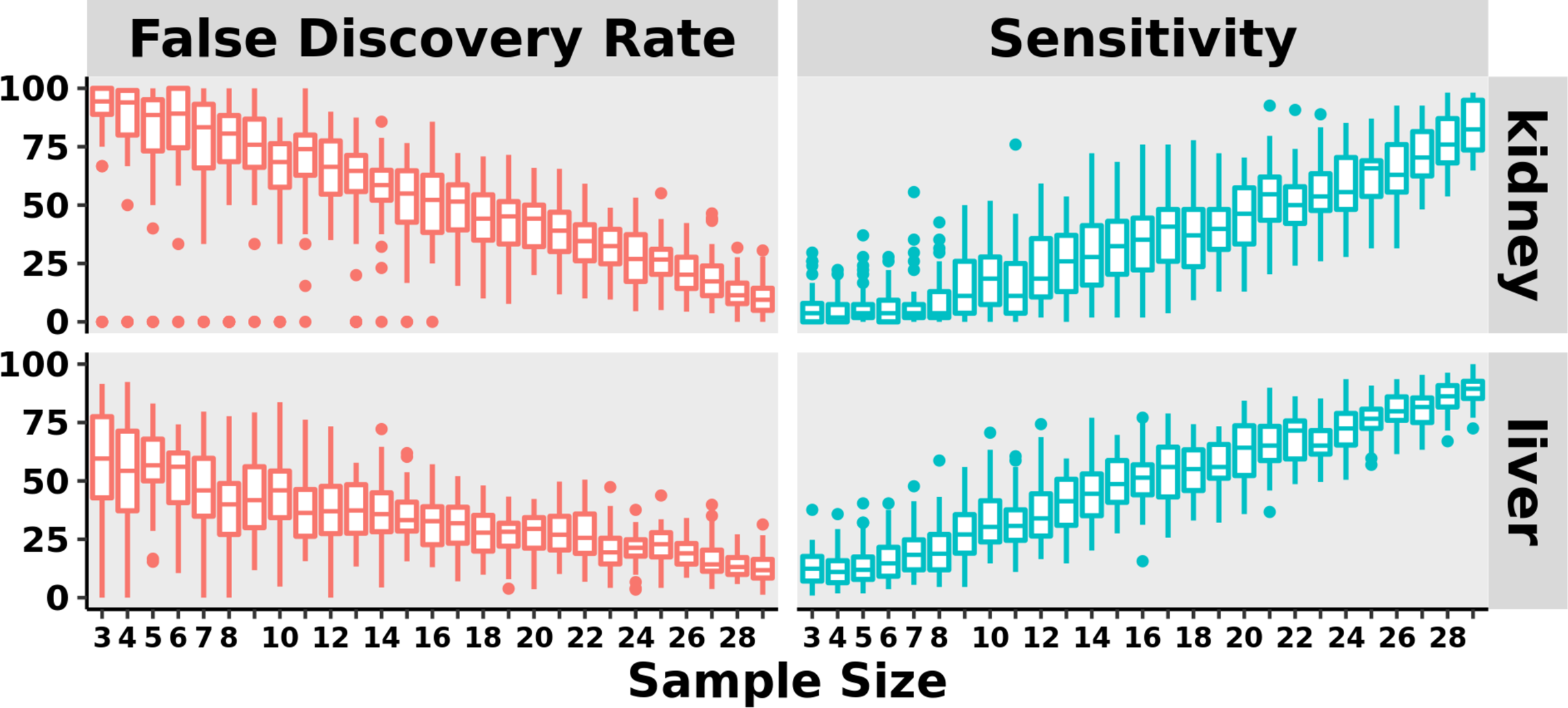
Impact of sample size on sensitivity & false discovery rate in Fat4 Hets. Plots are constructed as in Figure 2A, but for a different genetic knockout, Fat4. Only kidney and liver had sufficient number of DEGs for a meaningful analysis.

**Supplementary Figure 6A.**
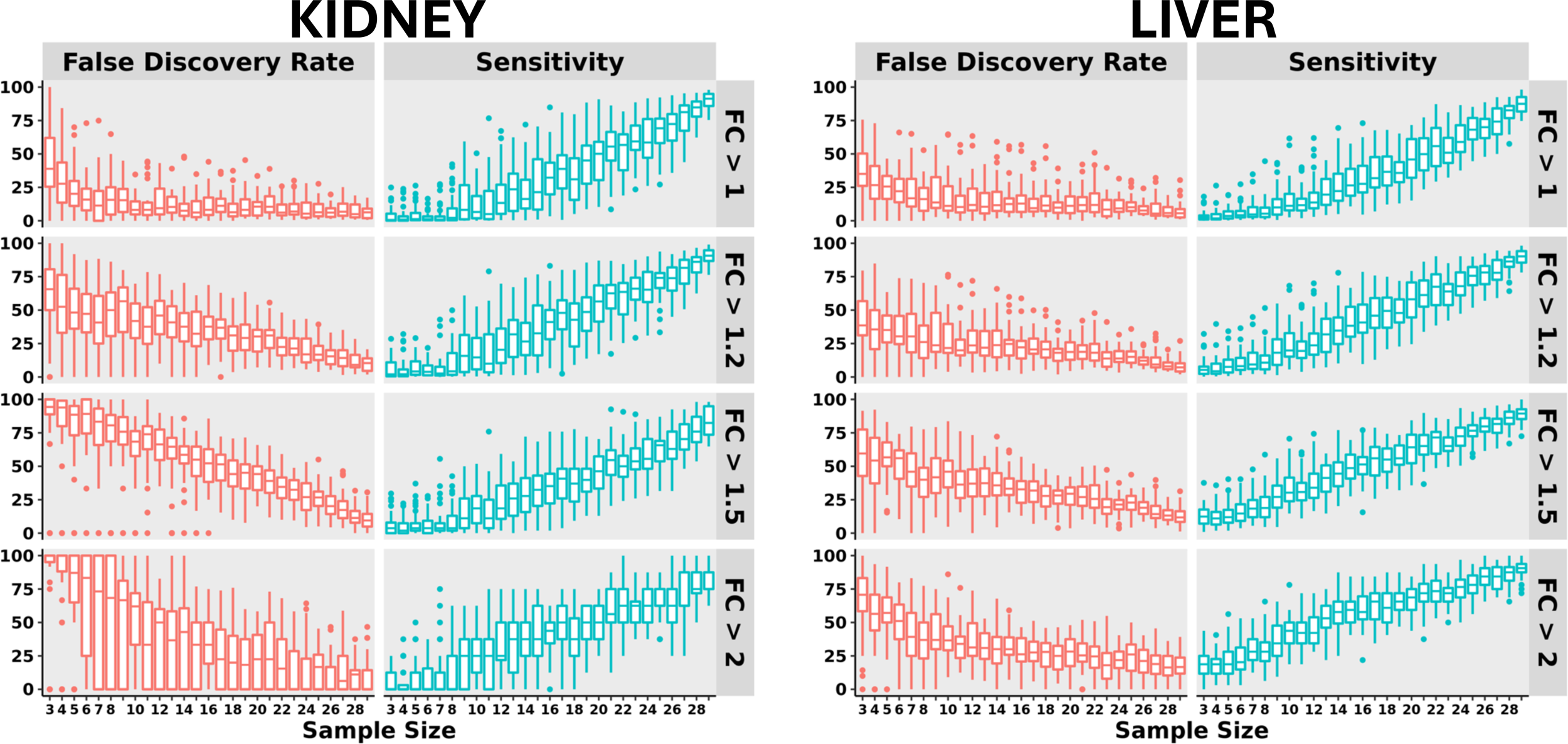
Impact of sample size and fold change threshold on sensitivity & false discovery rate in Fat4 Het mice. Plots are constructed as in Supplementary Figure 1A, but for a different genetic knockout, Fat4. Only kidney and liver had sufficient number of DEGs for a meaningful analysis.

**Supplementary Figure 6B.**
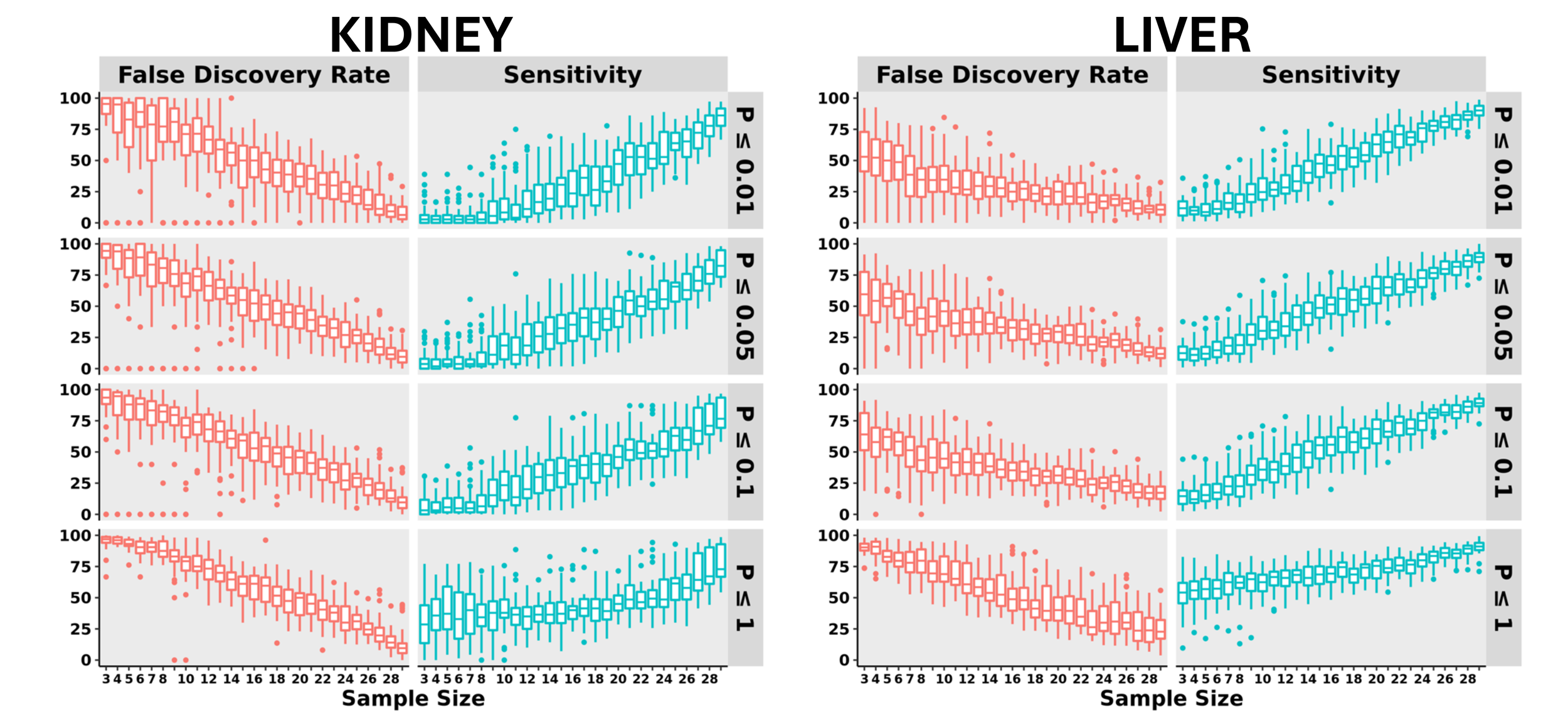
Impact of sample size and P-value threshold on sensitivity & false discovery rate in Fat4 Het mice. Plots are constructed as in Supplementary Figure 1B, but for a different genetic knockout, Fat4. Only kidney and liver had sufficient number of DEGs for a meaningful analysis.

**Supplementary Figure 7.**
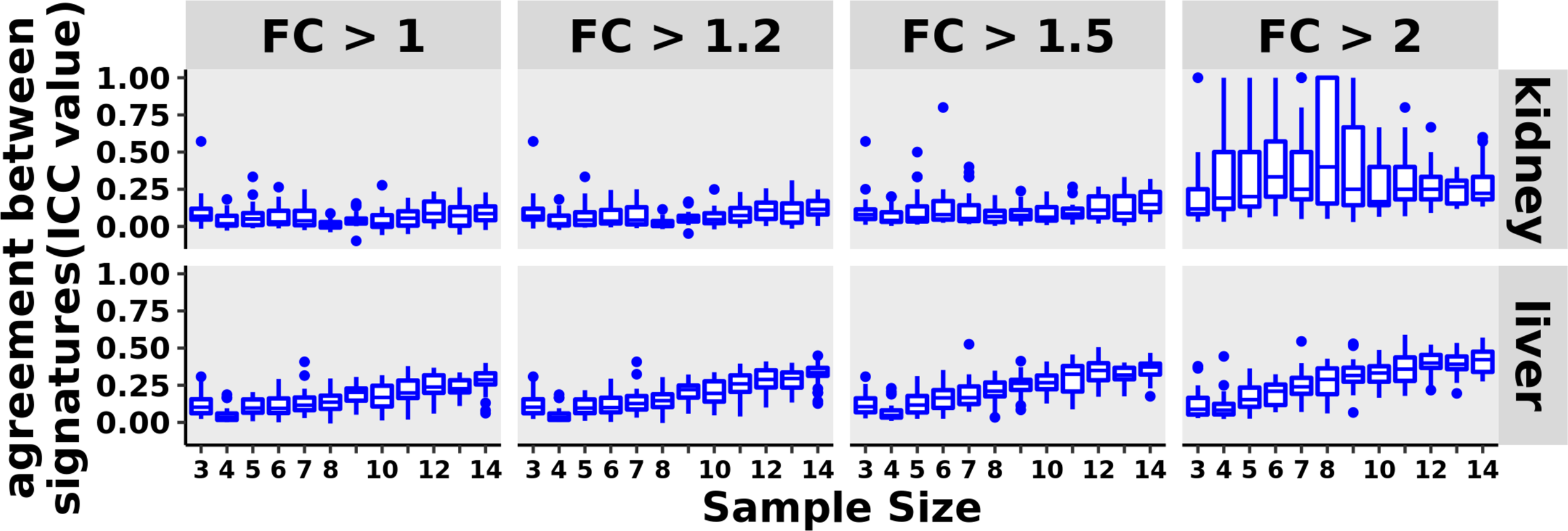
Agreement between DEGs obtained using disjoint method in Fat4 Het mice. Plots are constructed as in Figure 3, but for a different genetic knockout, Fat4. Only kidney and liver had sufficient number of DEGs for a meaningful analysis.

**Supplementary Figure 8.**
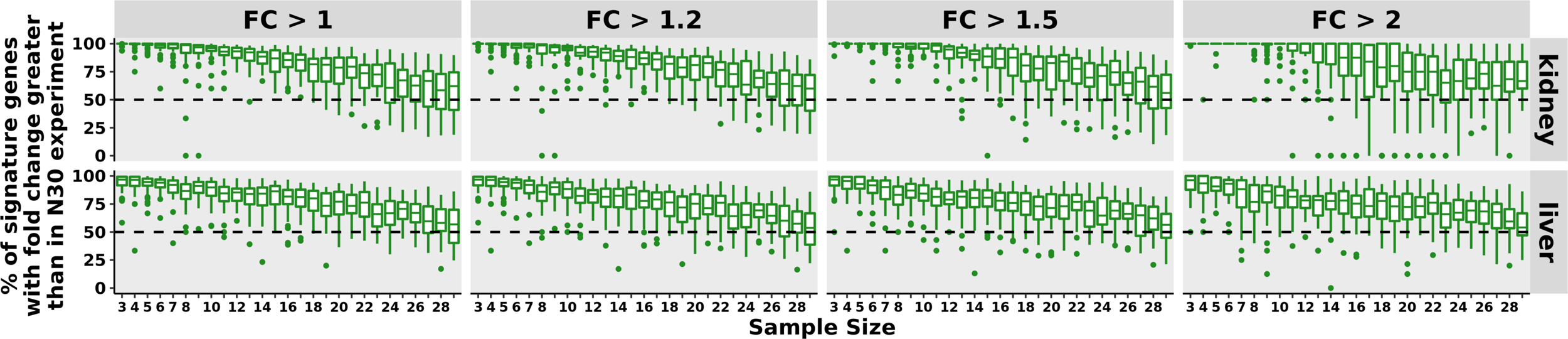
Impact of sample size on estimated effect size in Fat4 Hets. Plots are constructed as in Figure 4, but for a different genetic knockout, Fat4. Only kidney and liver had sufficient number of DEGs for a meaningful analysis.

